# From Viral Infections to Alzheimer’s Disease: Unveiling the Mechanistic Links Through Systems Bioinformatics

**DOI:** 10.1101/2023.12.05.570187

**Authors:** Anna Onisiforou, Panos Zanos

**Affiliations:** Translational Neuropharmacology Laboratory, Department of Psychology, University of Cyprus, Nicosia, 2109, Cyprus

**Keywords:** virus-host-disease interactions, Alzheimer’s disease, microbial pathogenesis, Herpes simplex virus 1, Human Cytomegalovirus, Epstein-Barr Virus, Hepatitis C Virus, SARS-CoV-2

## Abstract

**Background:** Emerging evidence suggests that certain microorganisms, including viral infections, may contribute to the onset and/or progression of Alzheimer’s Disease (AD), a neurodegenerative condition characterized by memory impairment and cognitive decline. However, the precise extent of their involvement and the underlying mechanisms through which specific viruses increase AD susceptibility risk remain elusive.

**Methods:** We used an integrative systems bioinformatics approach to identity viral-mediated pathogenic mechanisms by which specific viral species, namely Herpes Simplex Virus 1 (HSV-1), Human Cytomegalovirus (HCMV), Epstein-Barr Virus (EBV), Kaposi Sarcoma-associated Herpesvirus (KSHV), Hepatitis B Virus (HBV), Hepatitis C Virus (HCV), Influenza A virus (IAV) and Severe Acute Respiratory Syndrome Coronavirus 2 (SARS-CoV-2), could facilitate the pathogenesis of AD via virus-host protein-protein interactions (PPIs). We also sought to uncover potential synergistic pathogenic effects resulting from the reactivation of specific herpesviruses (HSV-1, HCMV and EBV) during acute SARS-CoV-2 infection, potentially increasing AD susceptibility.

**Results:** Our findings show that *Herpesviridae* Family members (HSV-1, EBV, KSHV, HCMV) impact AD-related processes like amyloid-beta formation, neuronal death, and autophagy. Hepatitis viruses (HBV, HCV) influence processes crucial for cellular homeostasis and dysfunction. Importantly, hepatitis viruses affect microglia activation via virus-host PPIs. Reactivation of HCMV during SARS-CoV-2 infection could potentially foster a lethal interplay of neurodegeneration, via synergistic pathogenic effects on AD-related processes like response to unfolded protein, regulation of autophagy, response to oxidative stress and amyloid-beta formation.

**Conclusions:** Collectively, these findings underscore the complex link between viral infections and AD development. Perturbations in AD-related processes by viruses can arise from both shared and distinct mechanisms among viral species in different categories, potentially influencing variations in AD susceptibility.

## Background

Alzheimer’s Disease (AD) is a progressive chronic neurodegenerative disease (ND) of the central nervous system characterized by memory impairment and cognitive decline [1]. The primary pathophysiological hallmarks of AD are the formation of amyloid plaques, neurofibrillary tangles, and neuronal loss in the brain [1]. Currently, there are no effective pharmacotherapies for its treatment [2]. It’s the predominant form of dementia in the elderly, affecting 55 million people worldwide [3]. AD is of multifactorial origin involving the complex interaction of both genetic and environmental risk factors [4]. Viral infections are environmental risk factors that have been hypothesized to increase susceptibility towards the development of AD [5]. Indeed, individuals seropositive for human Herpes Simplex Virus 1 (HSV-1) show increased risk of developing AD, and increased presence of HSV-1 DNA was found in AD post-mortem brains [6–8]. The detection of infectious agents, such as HSV-1, in the brain of AD patients led to the “antimicrobial protection hypothesis of AD”, which suggests that increased microbial burden in the brain might lead to the disposition of β-amyloid due to the role of Amyloid-beta (Aβ) in innate immune responses against pathogens, leading to sustain neuroinflammation that propagates neurodegeneration [9]. Several other viruses have been also linked to the development of AD, including Herpes Simplex Virus 2 (HSV-2), Human Cytomegalovirus (HCMV), Epstein-Barr Virus (EBV), Varicella-Zoster Virus (VZV), Human Herpesvirus 6A/B (HHV-6A/B), Human Herpesvirus 7 (HHV-7), Hepatitis C Virus (HCV), and severe acute respiratory syndrome coronavirus 2 (SARS-CoV-2) [6,10–15]. Overall, the current evidence suggests that viral infections may increase the risk of developing AD and/or facilitate its progression. However, further investigation is warranted to understand the viral-mediated pathogenic mechanisms through which different viral species contribute to AD susceptibility.

Co-infection with two viruses can have synergistic pathogenic effects, potentially increasing susceptibility to AD development. For instance, reactivation of latent viruses may occur during acute infection with respiratory viruses. Evidence suggests that SARS-CoV-2 infection can trigger the reactivation of latent herpesviruses like EBV [16], HCMV [17] and HSV-1 [18]. This reactivation may contribute to COVID-19 severity and post COVID-19 cognitive symptoms resembling AD, like memory impairment and brain fog [16,19–22]. Our previous work, leveraging systems bioinformatics methodologies, also suggested an elevated risk of AD development following SARS-CoV-2 infection [23]. Hence, it is not surprising that there was an observed approximately 17% increase in AD-related deaths in 2020, possibly associated with COVID-19 infections [24]. Thus, it is important to understand the mechanisms by which the synergistic action of two viruses could potentially increase the risk of AD.

Computational approaches, especially network-based methodologies, have been widely employed to shed light on microbe-host interactions underlying the onset of an array of diseases [25,26]. We have previously developed and applied innovative network-based approaches that allowed us to gain new insight on the role of existing viruses, but also emerging viruses (such as SARS-CoV-2), in the development of NDs [23,27,28]. In this study, we extend our previous research [23,27,28] to investigate viral-mediated pathogenic mechanisms that might lead to AD through virus-host protein-protein interactions (PPIs), with a focus on specific viral species (HSV-1, SARS-CoV-2, EBV, HCV, HCMV, Kaposi Sarcoma-associated Herpesvirus (KSHV), Hepatitis B Virus (HBV) and Influenza A virus (IAV)). **Figure 1** illustrates the main steps of our methodology. We also explore potential synergistic pathogenic effects that might arise from the reactivation of herpesviruses during SARS-CoV-2 infection, potentially increasing AD susceptibility.

## Methods

### Reconstruction and Analysis of the AD KEGG Pathway-Pathway Network

To identify pathways linked to AD development, our initial step involved utilizing the *String: disease app* within Cytoscape to retrieve the top 200 disease-associated proteins of AD (ID:10652), determined by their highest disease score. Subsequently, using the *ClueGO app* [29] within Cytoscape we performed enrichment analysis on these proteins, utilizing the Kyoto Encyclopedia of Genes and Genomes (KEGG) database. Specifically, we regarded pathways with an adjusted *p*-value of ≤ 0.01 (corrected with Bonferroni step-down) as statistically significant.

**Figure 1:**
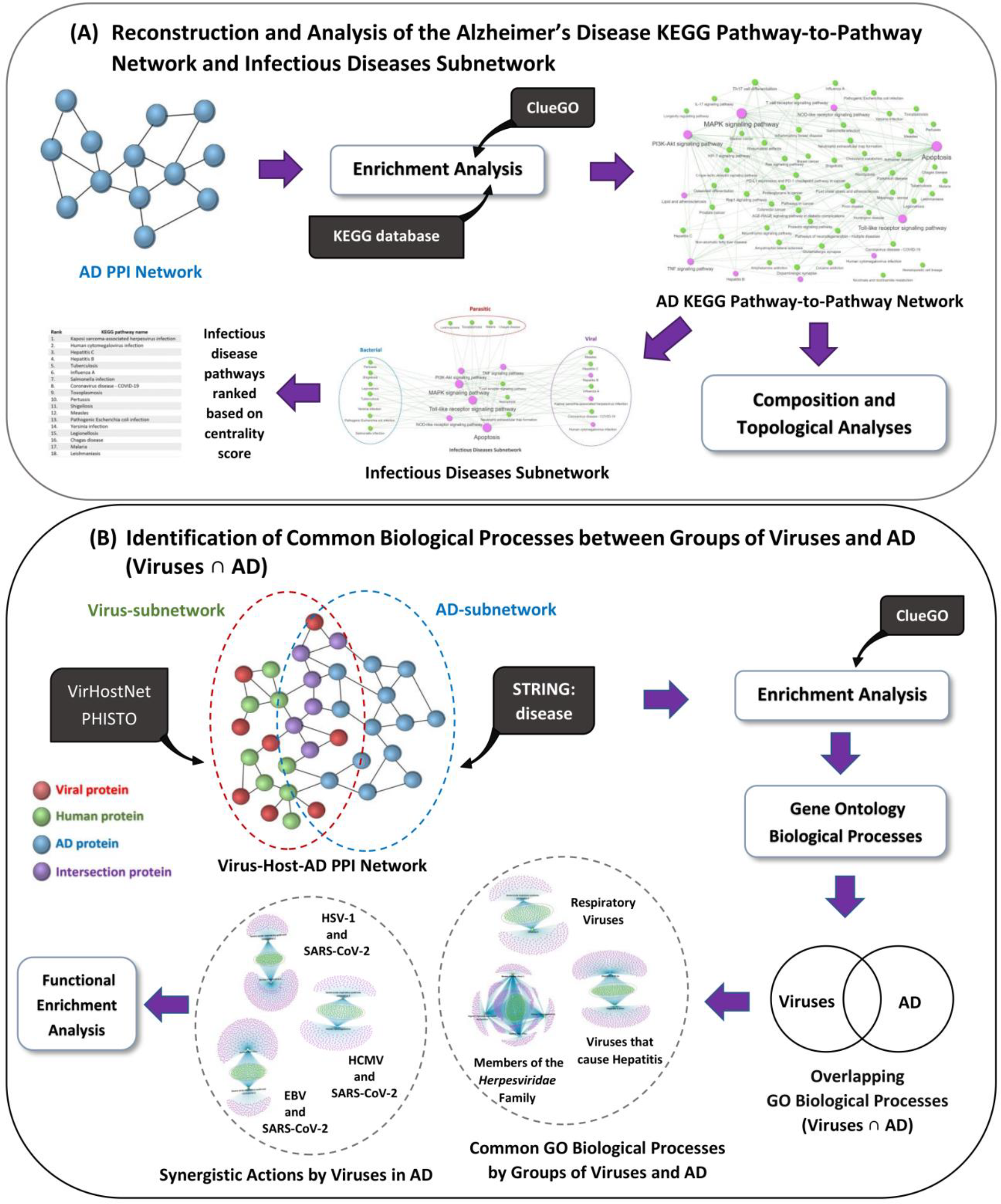
Schematic representation of our methodology implemented in this study. The aim of the present study was to identify potential pathogenic mechanisms by which specific viral species may lead to AD. **(A)** We reconstruct and analyze the AD KEGG pathway-to-pathway network. We also extract the Infectious Diseases subnetwork from the AD KEGG pathway network to investigate the effects of infectious-disease pathways in AD. **(B)** We present the data sources employed to reconstruct the Virus-host-AD PPI networks for each of the eight viral species (HSV-1, SARS-CoV-2, EBV, HCV, HCMV, KSHV, HBV, IAV) under investigation. We outline the methodology used to identify overlapping Gene Ontology (GO) biological processes (Viruses ∩ AD) between each viral species and AD via enrichment analysis. We also highlight common pathological mechanisms in AD by three groups of viral species: members of the Herpesviridae family (HSV-1, EBV, KSHV and HCMV), viruses that cause hepatitis (HCV and HBV) and respiratory viruses (SARS-CoV-2 and IAV). Lastly, we explore potential synergistic viral-pathogenic effects that could contribute to AD development through the reactivation of Herpesviruses (HCMV, HSV-1 and EBV) during acute SARS-CoV-2 infection.

Furthermore, we reconstructed the AD KEGG pathway-to-pathway interaction network. By utilizing the KEGGREST package [30] in R, we parsed each of the 353 KEGG pathways (Homo Sapiens) entries contained in the KEGG database [31] to extract information of the functional relationships of each pathway with other pathways. We then combined all the interactions obtained, which resulted into 1914 functional pathway-to-pathway interactions. Functional relationships between pathways, represent the interconnectedness and communication that occurs between different pathways to accomplish complex physiological processes, such as where components from one pathway influence the activity or regulation of components in another pathway. Subsequently, we extracted the functional relationships related to the 64 KEGG pathways found through enrichment analysis associated with the top 200 disease-associated proteins of AD. This resulted in constructing an AD pathway-pathway network with 64 nodes (pathways) and 177 edges (functional interactions).

Additionally, we conducted composition analysis on the AD KEGG pathways network to determine the subclasses to which the 64 AD pathways belong. To achieve this, we used the KEGGREST package [30] in R to extract the subclass classification of each of the 64 pathways based on their classification in the KEGG database.

Furthermore, by using the igraph package in R [32], we performed topological analysis on the AD KEGG pathways network to evaluate the importance of the infection-related pathways in interacting with and influencing key AD pathways. This was achieved by measuring their centrality score within the network, calculated by summing five topological measures: hub score, degree, betweenness, closeness and eigenvector centrality. The infectious diseases pathways were then ranked based on highest to lowest centrality score, with a high score indicating a high influence on AD.

### Reconstruction of Integrated Virus-Host-AD PPI Networks

We initially conducted a thorough literature review analysis to identify viruses associated with the development of AD. Our investigation led us as to identify 10 specific viruses that are linked with AD, namely HSV-1, HSV-2, HCMV, EBV, VZV, HHV-6A/B, HHV-7, HCV, SARS-CoV-2 [6,10–15]. Next, we collected experimentally validated virus-host PPIs from the PHISTO [33] and VirHostNet 3.0 [34] databases (date 1 August 2023). We also obtained virus-host PPIs for IAV, HBV and KSHV. These viruses were found through the analysis of the AD KEGG pathway-pathway network as having the highest centrality score in influencing other AD-related pathways. Out of the thirteen viruses, virus-host PPIs were available for twelve of them. Four viruses were excluded due to having less than 100 virus-host PPIs available. Duplicated entries from the two databases were eliminated. Table 1 provides detailed characteristics of the eight viruses and the number of virus-host PPIs collected for further analysis from each viral species.

**Table 1:**
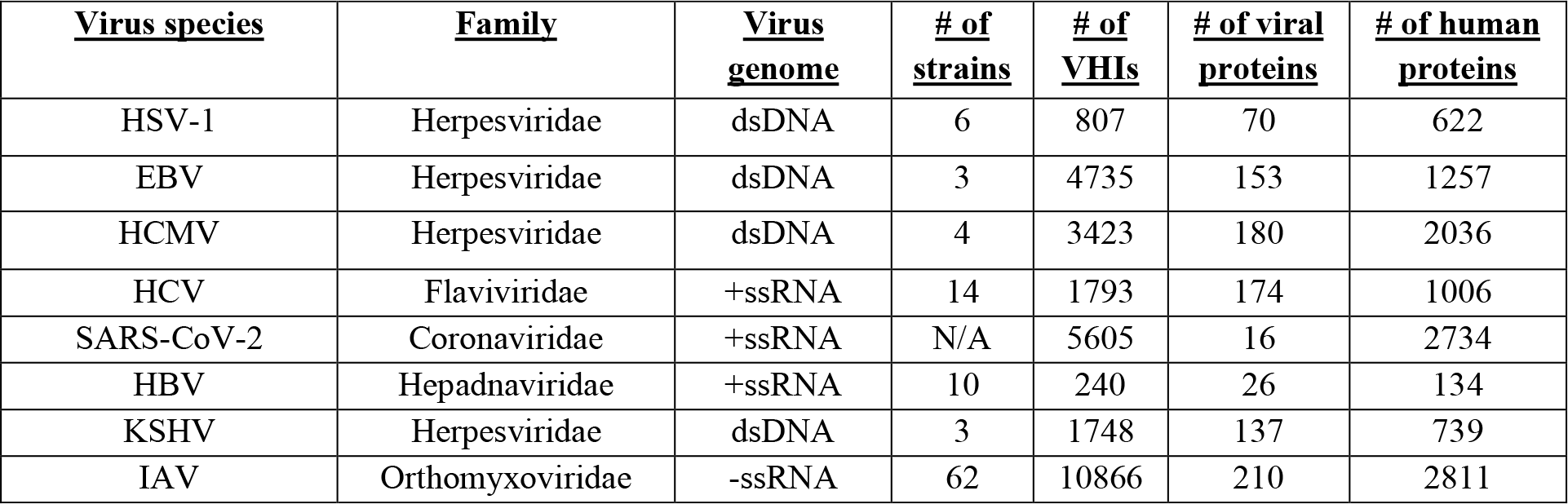
Characteristics of the Virus - Human host PPI networks.

We then reconstructed eight integrated virus-host-AD PPI networks (see **Supplementary File 1: Table 1**). to identify the viral-mediated pathogenic mechanisms through which the eight viral species (**Table 1**) could potentially contribute to the development and/or progression of AD. To generate the integrated networks, we merged the virus-host PPI network of each viral species with the AD PPI network, which consisted of the top 200 disease-associated proteins of AD. The confidence cut-off score, involving the interactions between the human proteins was set at 0.8, as in our previous works [23,28]. The confidence score is determined based on the nature and quality of the supporting evidence for the PPIs, with values ranging from 0 (indicating low confidence) to 1.0 (indicating high confidence). As the score increases, the likelihood of the PPIs being true positives also rises [35]. Consequently, a more rigorous cut-off of 0.8 was selected with the aim of improving reliability and minimizing the inclusion of false positive results.

### Identification of Common Gene Ontology Biological Processes between Groups of Viruses and AD (Viruses ∩ AD)

To identify common biological processes among the eight viral species (HSV-1, SARS-CoV-2, EBV, HCV, HCMV, KSHV, HBV, IAV) through which they might contribute to AD, we classified them into three groups based on their type and disease they cause: (a) Members of the *Herpesviridae* Family (HSV-1, EBV, KSHV and HCMV); these viruses share genetic and structural similarities, are known for their ability to establish latent infections in their hosts and most display a high degree of neurotropism, (b) Viruses that Cause Hepatitis (HCV and HBV); target the liver and can lead to various degrees of liver damage and inflammation, (c) Respiratory Viruses (SARS-CoV-2 and IAV); primarily target the respiratory tract and cause respiratory illness.

We then employed a methodology applied in our previous work [23], which first involves isolating two subnetworks from each of the eight constructed integrated virus-host-AD PPIs networks. For each integrated network we extracted the (i) virus subnetwork, which includes the human protein targets of each viral species proteins and their first neighbors and (ii) the AD-related subnetwork, which includes the 200 disease-associated proteins of AD and their first neighbors.

Then, we performed enrichment analysis on the human proteins contained in each of the extracted subnetworks (see **Supplementary File 1: Table 1**) by using the Gene Ontology Biological Processes (GO BP) database and retaining only significant processes that had an adjusted *p*-value ≤0.001 (corrected with Bonferroni step-down). For the enrichment analysis of the extracted subnetworks, we used a stricter *p-* value, than in enrichment analysis of the disease-associated proteins of AD, because the integrated virus-host-AD PPI networks contain a significant larger number of human proteins than the AD PPI network. Subsequently, Venn diagrams were used to identify the overlapping GO BP between the enriched results of the AD subnetwork and viral subnetwork associated with each of the eight integrated virus-host-AD PPI networks (see **Supplementary File 1: Table 2**).

Finally, for each group of viruses (Members of the *Herpesviridae* Family, Viruses that Cause Hepatitis, Respiratory Viruses) we created GO BP-Viruses networks. These networks included the overlapping GO BP (Viruses ∩ AD) that were identified as being modulated by each viral species in AD. Subsequently, we identified the common GO biological processes that could potentially be influenced by all viruses contained in each group. We then performed functional enrichment analysis on the common GO BP to determine the functional groups that these GO BP belong to.

### Potential Synergistic Viral-mediated Pathogenic Effects in AD

To explore the potential for synergistic viral-mediated pathogenic effects in AD, we also identified common GO BP influenced by the reactivation of herpesviruses (HSV-1, HCMV and EBV) during acute SARS-CoV-2 infection. Using the overlapping GO BP (Viruses ∩ AD), we constructed three GO BP-Viruses networks (SARS-CoV-2 and EBV, SARS-CoV-2 and HSV-1, SARS-CoV-2 and HCMV) and performed functional enrichment analysis on the common GO BP.

## Results

### Infectious Diseases Pathways and their Role in AD

We reconstructed and analyzed the AD KEGG pathway-pathway network (see **Figure 2A**), consisting of 64 nodes (pathways) and 177 edges (interactions) to explore the potential involvement of viral infections in AD. This network was constructed using the 64 statistically significant pathways found through enrichment analysis, to be linked with the top 200 disease-related proteins associated to AD from various sources.

**Figure 2:**
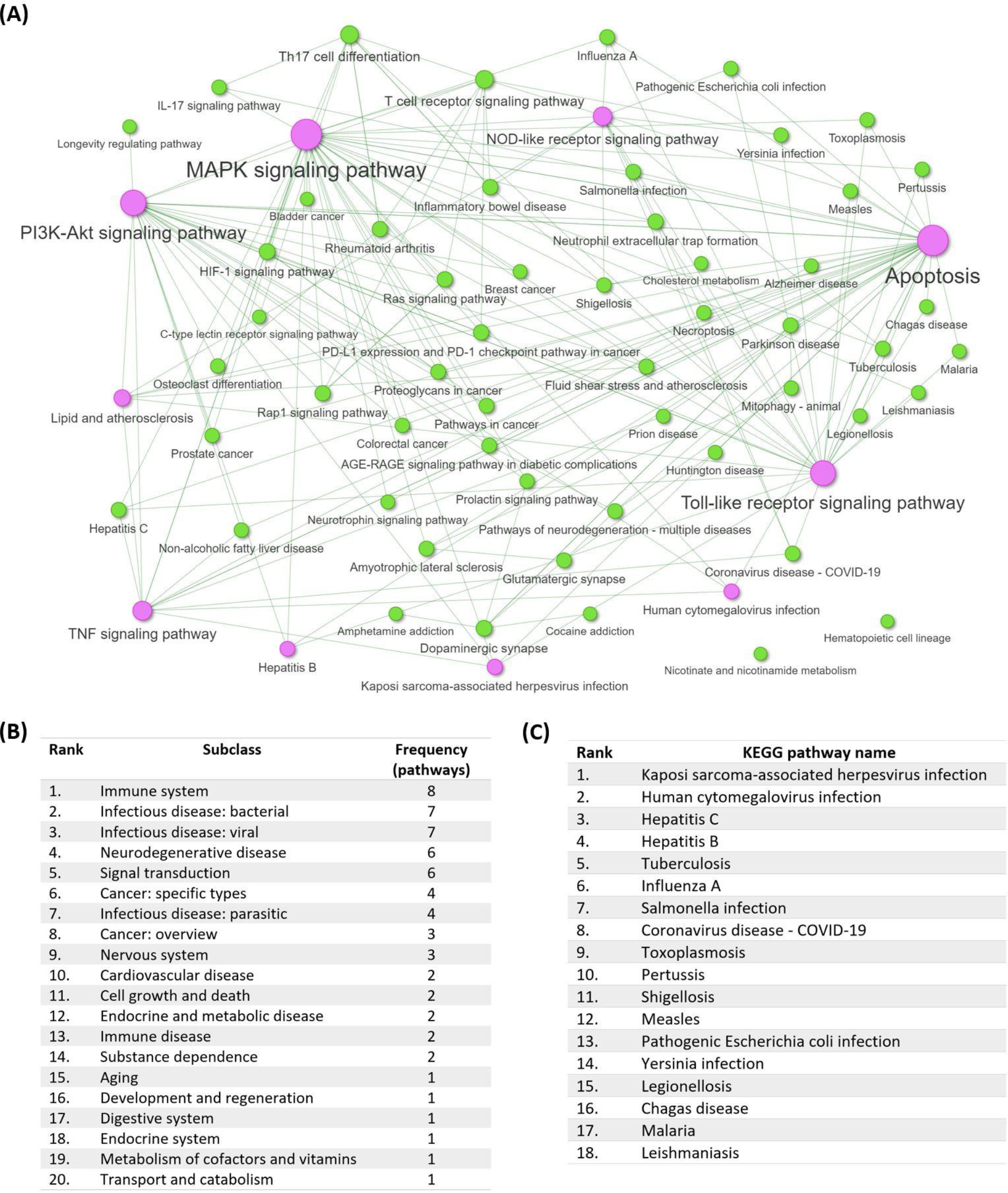
Impact of Infectious diseases pathways in AD pathophysiology. **(A)** Visualization of the AD KEGG pathway-pathway network, indicating the functional relationships between the 64 AD-related pathways. The top 10 hubs within the network are highlighted in purple color. The size of each node is proportional to its degree, reflecting the number of connected edges for that specific node. **(B)** Top 10 subclasses found in the AD KEGG pathway-pathway network and the number of pathways found in each subclass. **(C)** Infection-related pathways ranked from highest-to-lowest centrality score in influencing other AD-related pathways in the network.

Composition analysis of the 64 pathways within this network revealed they belong to 20 subclasses (see **Figure 2B**) according to the KEGG database classification system. Among them, 18 pathways were related to infectious diseases, with seven associated with viral infections, seven with bacterial infections, and four with parasitic infections. This suggests a potential link between these infectious diseases and the pathogenesis of AD, as these pathways contained more overrepresented genes related to AD than would be expected by chance. This means that these infectious diseases modulate or influence genes associate with AD.

Furthermore, using topological analysis, we assessed the significance of each of the 18 infection-related pathways within the AD KEGG pathway-pathway network regarding their influence on other AD key pathways. We determined the significance of each infection-related pathway by calculating its centrality within the network, using a combined score based on five topological measures (hub score, degree, betweenness, closeness, and eigenvector centrality). Higher centrality scores indicate pathways crucial for information flow and control among network pathways. Consequently, high centrality score suggests greater capability of the infection-related pathway to impact other AD-related pathways. This a critical aspect to consider when comprehending the intricate interplay between infections and AD pathogenesis. Notably, the analysis revealed that members of the *Herpesviridae* family (including KSHV and HCMV) as well as members of the *Flaviviridae* family (HCV), and *Hepadnaviridae* family (HBV) had the highest centrality scores in influencing other AD-related pathways (**Figure 2C**).

To further investigate the influence of infectious disease pathways in AD, we extracted the infectious diseases subnetwork (see **Figure 3**) from the AD KEGG pathways network. This analysis revealed that the 18 infection-related pathways functionally interact with 9 other AD pathways, of which 6 act as hubs within the network. Interaction with these hub nodes may enable pathogens to facilitate the development of AD through systemic pathogenic effects.

**Figure 3:**
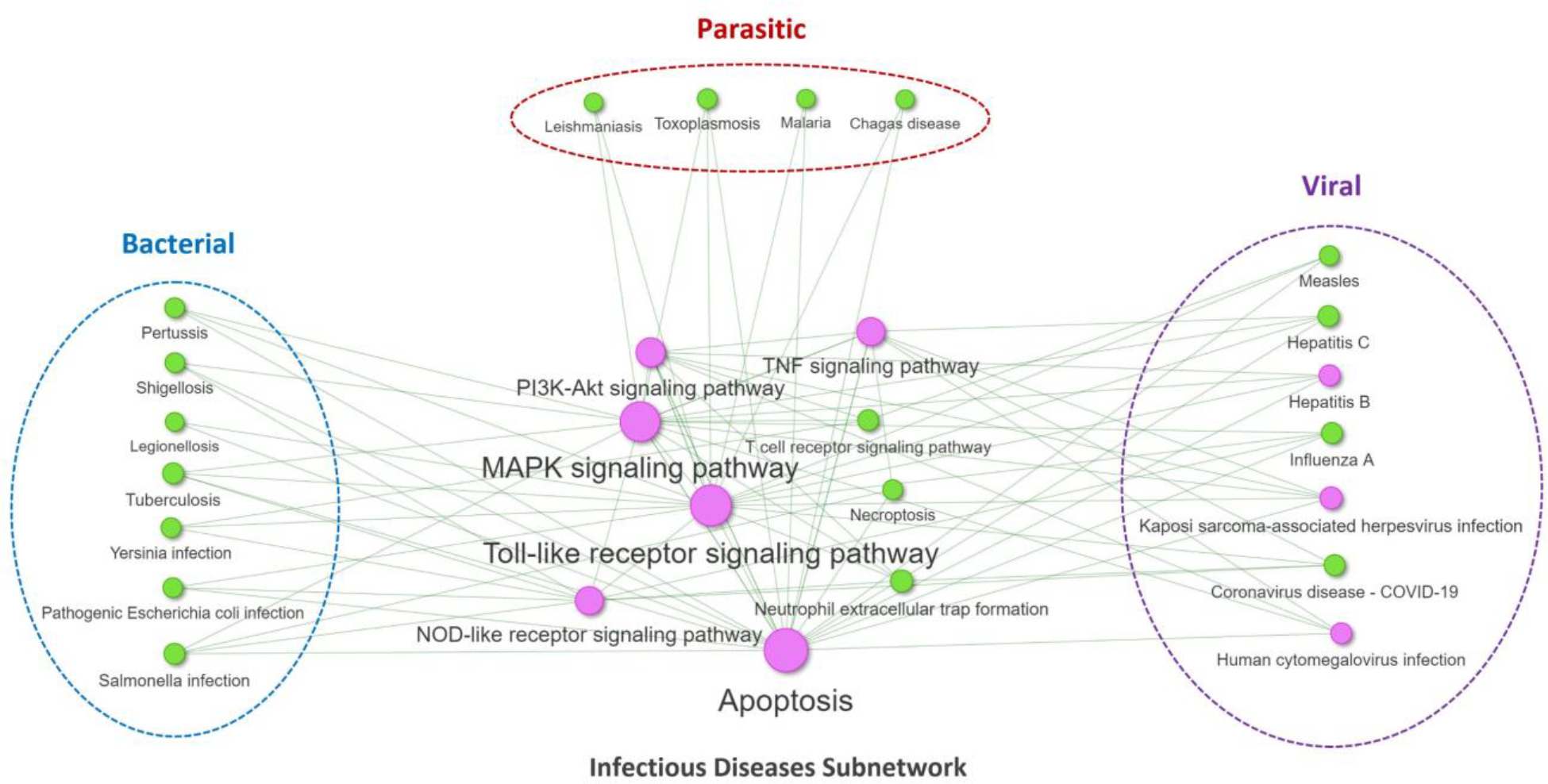
Visualization of the Infectious Diseases Subnetwork isolated from the AD KEGG pathway-pathway network. We highlight the 18 infection-related pathways, grouped based on their subclass (viral, bacterial, parasitic), and indicate their interaction with their first neighbors’ pathways within the AD KEGG pathways network. The first neighbors’ pathways include 6 out of the 10 top nodes in the network that act as hubs (highlighted in purple).

### Common Viral-mediated Pathogenic Mechanisms by Groups of Viruses in AD

To identify common viral-mediated pathogenic mechanisms among groups of viruses in AD we categorized the eight viral species into three groups: (a) Members of the *Herpesviridae* Family (HSV-1, EBV, KSHV and HCMV), (b) Viruses that Cause Hepatitis (HCV and HBV), (c) Respiratory Viruses (SARS-CoV-2 and IAV). Integrated GO BP-Viruses networks were constructed for each group (**Figure 4A, C, E**) using the overlapping GO BP (Viruses ∩ AD) identified for each virus with AD. Functional enrichment analysis was performed to identify the functional groups these common processes belong to (**Figure 4B, D, F**).

**Figure 4:**
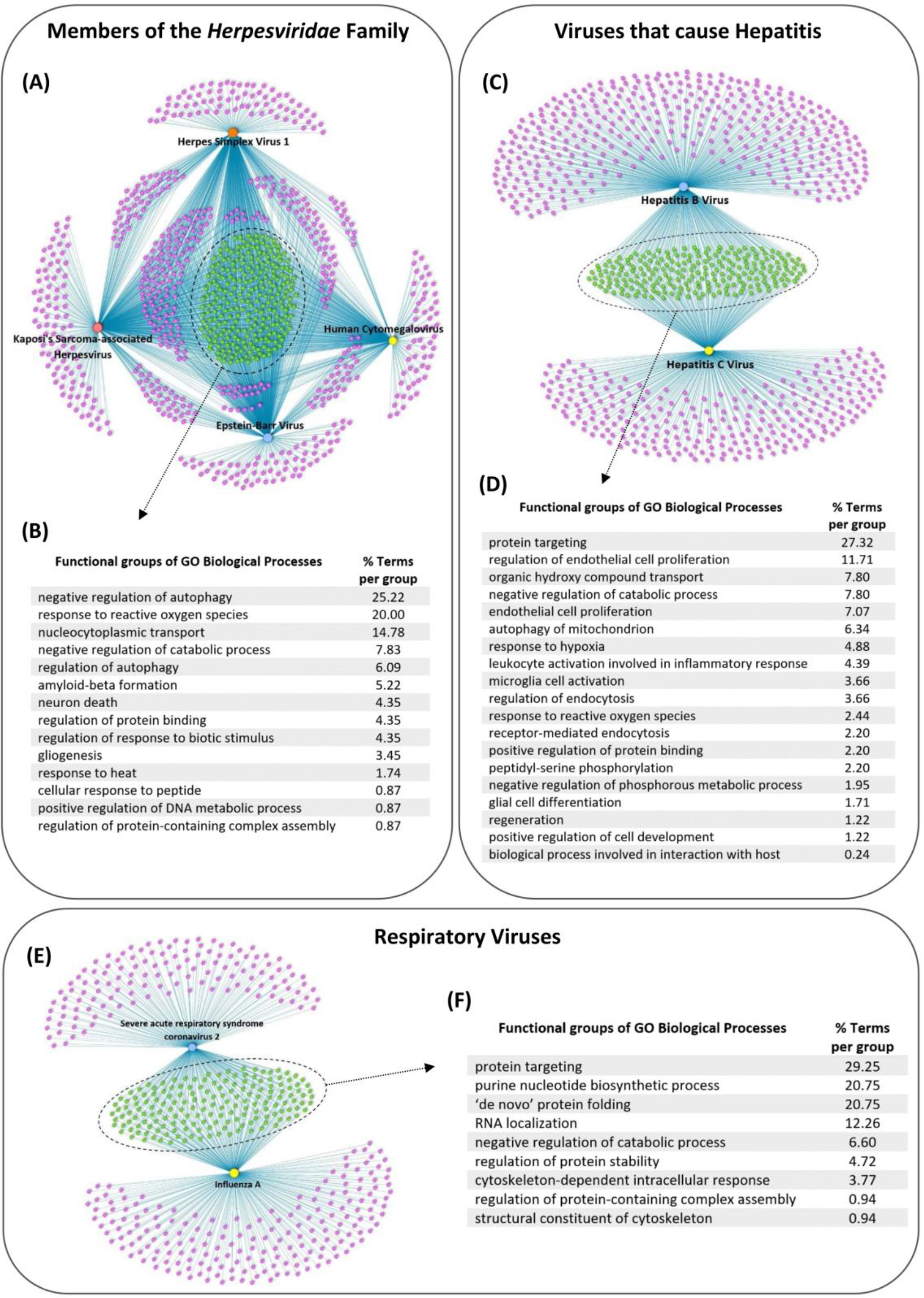
Common viral-mediated pathogenic mechanisms by which groups of viruses can potentially contribute to the development of AD. Integrated GO BP-Viruses networks (**A, C, E**) containing the overlapping GO BP (Viruses ∩ AD), found for each virus with AD for (i) Members of the Herpesviridae Family (HSV-1, EBV, KSHV and HCMV), (ii) Viruses that Cause Hepatitis (HCV and HBV) and (iii) Respiratory Viruses (SARS-CoV-2 and IAV). Functional groups to which the common GO BP of each network are associated (**B, D, F**) as determined through functional enrichment analysis.

The integrated network encompassing the four Members of the *Herpesviridae* Family (HSV-1, EBV, KSHV and HCMV) unveiled a total of 241 common GO BP, belonging to 14 functional groups, by which they can facilitate the development of AD (**Figure 4A, B**). Notably, 20.00% of the terms belonging to the group of “response to reactive oxygen species” (ROS). Additionally, 25.22% to “negative regulation of autophagy” and 6.09% to “regulation of autophagy”. Furthermore, 5.22% fall within “amyloid-beta formation” group, 4.35% to “neuron death” and 3.45% to “gliogenesis” group. These findings collectively suggest that herpesviruses have the capacity to impact diverse biological processes associated with key pathological features of AD, indicating their potential role in AD pathogenesis.

Furthermore, the integrated network containing the two viruses that cause hepatitis (HCV and HBV) revealed a total of 219 common GO BP that belong to 19 functional groups by which they can lead to AD (**Figure 4C, D**). Nearly one-third of the terms, 27.32%, belong to the “protein targeting” functional group, which involves the transport and precise localization of proteins to specific cellular regions, including the peroxisome and the mitochondrion. Disruptions can lead to protein mislocalization, which could potentially contribute to the accumulation of Aβ. Moreover, a considerable number of terms are linked to processes directly associated with endocytosis, with 3.66% of terms associated with “regulation of endocytosis” and 2.20% to the “receptor-mediated endocytosis” group. Furthermore, 6.34% of terms are allocated to the “autophagy to mitochondrion” classification, 4.88% to “response to hypoxia” and 2.44% to “response to reactive oxygen species”. Notably, certain terms find their place within functional groups closely associated with inflammation, with 3.66% associated with the “microglia cell activation” and 4.39% in the “leukocyte activation involved in inflammatory response”.

Moreover, the integrated network encompassing the two respiratory viruses (SARS-CoV-2 and IAV) unveiled a total of 172 common GO BP that belong to 9 functional groups (see **Figure 4E, F**). Many of these terms are related to protein processes, with 29.25% in “protein targeting,” 20.75% in “de novo protein folding”, 4.72% in “regulation of protein stability”, and 0.94% in “regulation of protein-containing complex assembly” functional group.

### Synergistic Actions between Herpesviruses and SARS-CoV-2 in the Pathogenesis of AD

During acute SARS-CoV-2 infection, herpesvirus reactivation may synergistically enhance viral-mediated pathogenic effects through virus-host PPIs, potentially increasing AD susceptibility risk and contributing to post COVID-19 AD-like cognitive symptoms. To determine the biological processes whose dysregulation might be amplified by the reactivation of each of the herpesviruses EBV, HSV-1, and HCMV during SARS-CoV-2 infection, we reconstructed three integrated GO BP-Viruses networks. The integrated GO BP-Viruses network involving EBV and SARS-CoV-2 revealed 186 common GO BP (**Figure 5A**) that belong to 8 functional groups (**Figure 5B**). Notably, a significant 70.21% of these processes are categorized under “protein targeting”. Additionally, 10.64% of terms involve “negative regulation of catabolic processes,” encompassing mechanisms that downregulate the breakdown of various substances, including cellular, lipid, protein, and glycolytic catabolic processes.

**Figure 5:**
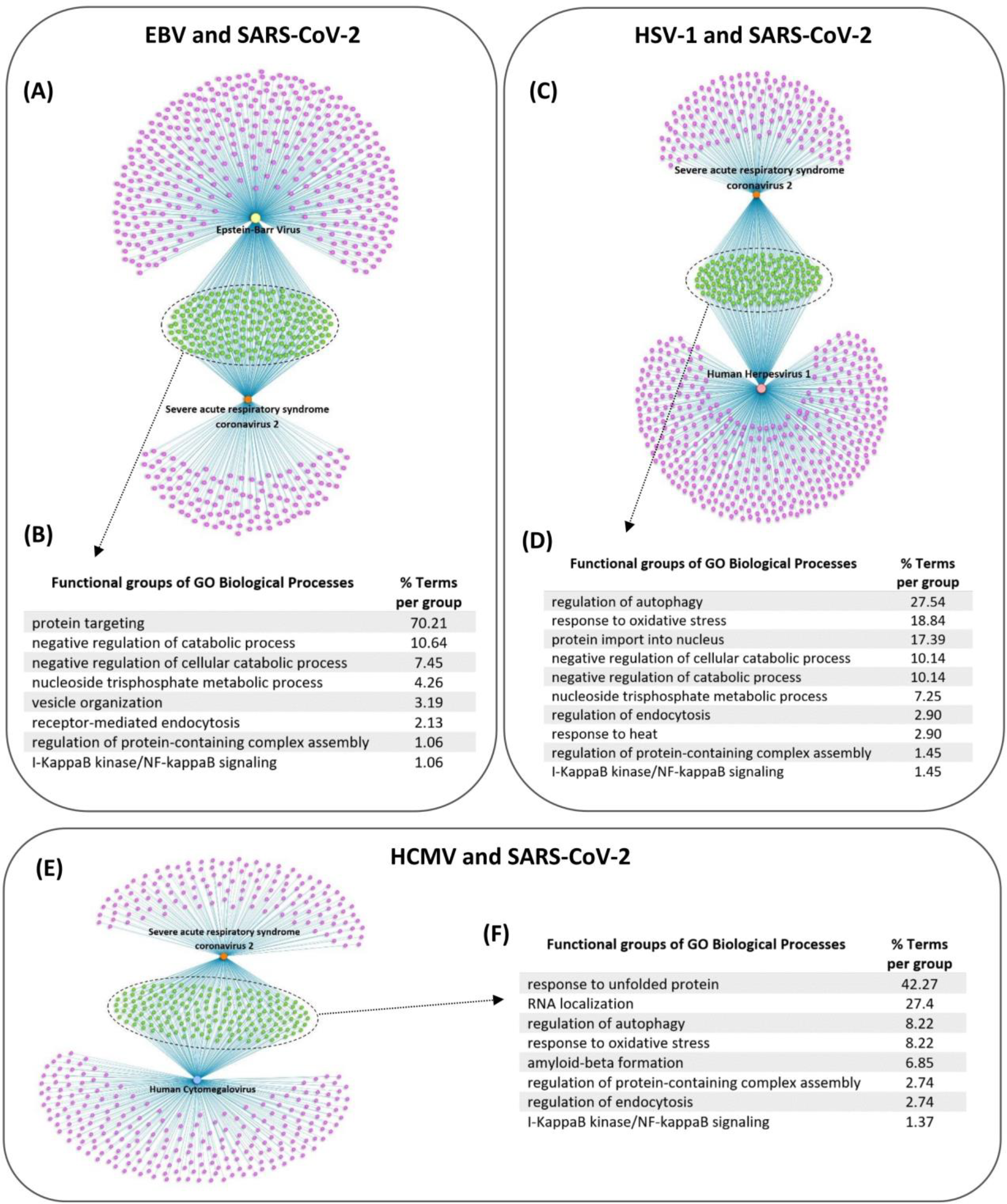
Synergistic Pathogenic Effects Between Herpesviruses and SARS-CoV-2 in AD Pathogenesis. Integrated GO BP-Viruses Networks and the Functional Groups of the common GO-BP: **(A-B)** EBV with SARS-CoV-2, **(C-D)** HSV-1 with SARS-CoV-2, and **(E-F)** HCMV with SARS-CoV-2.

Moreover, the integrated GO BP-Viruses network involving HSV-1 and SARS-CoV-2 revealed 152 common GO BP by which they can exert synergistic pathogenic effects, contributing to AD (**Figure 5C**). These processes are categorized into 10 functional groups (**Figure 5D**), with about one-third of the terms (27.54%) associated with the “regulation of autophagy”. Furthermore, 18.84% of the terms are associated with the “response to oxidative stress” group, and a significant 20.28% are linked to functional groups involved in the negative regulation of catabolic processes.

Lastly, the integrated GO BP-Viruses network involving HCMV and SARS-CoV-2 revealed collective 169 common GO BP, which are classified into 8 functional groups (**Figure 5E, F**). Of notable significance, approximately 42.27% of the terms are associated with “response to unfolded protein”, referring to events that leads to changes in the state or behavior of a cell or organism in reaction to a stimulus arising from a misfolded protein. Furthermore, they have the potential to exert synergistic actions via processes related to “regulation of autophagy” (8.22%), “response to oxidative stress” (8.22%), “amyloid beta formation” (6.85%) and “regulation of endocytosis” (2.74%).

## Discussion

Using an integrative network-based systems bioinformatics approach, we identified specific viral-mediated mechanisms by which eight viral species (HSV-1, SARS-CoV-2, EBV, HCV, HCMV, KSHV, HBV and IAV) could contribute to the development and/or progression of AD. We also isolated common viral-mediated pathogenic mechanisms that can lead to AD among groups of viral species. Additionally, we highlighted the potential for synergistic pathogenic effects during co-infection with two viruses, specifically the reactivation of herpesviruses (HSV-1, EBV and HCMV) during SARS-CoV-2 infection. This synergy can amplify disruptions in AD-related processes via virus-host PPIs, potentially increasing susceptibility to AD.

Herpesviruses, particularly HSV-1, are associated with increased risk in developing AD [6–8,36,37]. Our analysis results indicate that members of the *Herpesviridae* Family (HSV-1, EBV, KSHV, and HCMV) have the capacity to influence several of the primary pathological hallmarks associated with AD through virus-host PPIs. These encompass processes such as the generation of ROS, formation of Aβ, neuronal cell death, gliogenesis, and autophagy [38–40]. Increased production of ROS and dysregulation of autophagy are common pathological mechanisms found in many NDs, including AD [41,42]. Additionally, Aβ accumulation stands as a pivotal hallmark of AD [40], and the potential influence of herpesviruses on this process raises the possibility that they could have a crucial role in the initiation or progression of AD.

The “Antimicrobial Protection Hypothesis of AD” proposes that Aβ could serve as a defense mechanism against brain infections, given its demonstrated antimicrobial effects against various pathogens, including viruses [9]. Importantly, there is evidence suggesting that Aβ oligomers can sequester herpesviruses within insoluble amyloid deposits [43]. Our current findings also indicate that herpesviruses have the capacity to influence processes involved in the formation of Aβ via virus-host PPIs. Through the evolutionary process, viruses have acquired various functions, including mimicking and interfering immune responses [44]. Thus, it is possible that herpesviruses interfere with the Aβ processes as a means to evade its antimicrobial properties and escape the immune system. This interaction could potentially lead to the perturbation of the host’s antiviral response through Aβ, resulting in a detrimental interplay that accelerates the deposition of Aβ, contributing to AD pathogenesis. Therefore, understanding the complex interplay between the immune system, herpesviruses, and Aβ is imperative in unraveling the mechanisms through which these interactions can foster Aβ plaque formation.

Cognitive impairment has been reported in chronic infection with hepatitis viruses, HBV and HCV; however, conflicting results exist whether they influence AD pathogenesis [45,46]. A recent investigation revealed that administering direct-acting antivirals for the treatment of HCV infection substantially diminishes the risk of mortality among patients afflicted with AD and associated dementia [47].Our results indicate that hepatitis viruses (HBV and HCV) can modulate a diverse range of processes crucial for cellular homeostasis and dysfunction including the regulation of protein targeting, endocytosis, response to hypoxia, ROS and autophagy in the mitochondrion. Importantly, our results show that they can influence microglia activation via virus-host PPIs. Consistent with our results, it was previously shown that the mouse hepatitis virus can directly infect and subsequently activate microglia in a virus-induced mouse model of human ND [48]. Additionally, evidence indicates that microglia activation correlates with cognitive dysfunction in chronic HCV infection [49]. Understanding the intricate interactions between hepatis virus and microglia activation could potentially open avenues for targeted therapeutic interventions to mitigate the impact of these viruses on neurodegeneration and cognitive decline.

The role of members of the Herpesviridae family (KSHV and HCMV) and hepatitis viruses (HCV and HBV) in the development/progression of AD is also highlighted through the reconstruction and topological analysis of the AD KEGG pathway-to-pathway network. These viruses exhibited the highest centrality scores in influencing other AD-related pathways within the network, including several pathways that act as hubs. Pathogens often target high centrality nodes, like hub nodes, which allows them to exert systemic pathogenic effects within the human host [50,51]. Thus, our results suggest that these viruses may pose a higher risk of AD susceptibility as they have the ability to affect multiple other AD-related pathways, including hub nodes, hence facilitating the development of AD through systemic pathogenic effects.

Virus-virus interactions (VVIs) are very common in nature and have the potential to modify the trajectory of an ongoing infection, often influencing the severity of viral diseases [52]. Co-infection with two viruses may lead to synergistic pathogenic effects, where this collaborative impact results in the amplification of the dysregulation of biological processes beyond what each virus could achieve independently. This interaction may intensify the disruption of host biological processes associated with the pathogenesis of AD, potentially leading to more severe pathological outcomes and an increased risk of AD susceptibility. Given the high link of both members of the *Herpesviridae* family and SARS-CoV-2 with the development and/or progression of AD, as well as the demonstrated capability of SARS-CoV-2 to lead to reactivation of these herpesviruses [16–18], we sought to isolate the specific biological processes that would be amplified from the reactivation of each specific herpesviruses (HSV-1, HCMV, and EBV) during acute SARS-CoV-2 infection.

Our findings revealed a significant shared impact of EBV and SARS-CoV-2 on biological processes related to protein targeting. Disruption of protein targeting, especially in neuronal cells, can result in protein mislocalization [53], which could potentially contribute to the accumulation of Aβ. Dysfunction in neuronal cells can also potentially trigger neuroinflammation, an established pathological feature of AD. Based on these results, co-infection with these two viruses could potentially lead to more pronounced disturbances in protein localization, thereby potentially exacerbating the accumulation of mis-localized or misfolded proteins, and contributing to the molecular pathology of AD.

Furthermore, our results also indicate that HSV-1 and SARS-CoV-2 share the ability to impact processes whose dysregulation is associated with AD, including autophagy and response to oxidative stress. In the context of AD, oxidative stress triggers neuronal cell death by prompting the autophagy of accumulated Aβ, which in turn causes the permeabilization of the lysosomal membrane, ultimately resulting in neuron death [54]. Thus, the ability of co-infection with HSV-1 and SARS-CoV-2 to influence both of these crucial processes implicated in AD pathogenesis can increase susceptibility risk for its development.

Similar to co-infection with HSV-1 and SARS-CoV-2, the combination of HCMV and SARS-CoV-2 co-infection also appears to have the potential to lead to amplify pathogenic effects on processes involving autophagy and response to oxidative stress. Furthermore, both HCMV and SARS-CoV-2 have the capacity to impact processes related to response to unfolded protein and Aβ formation. These shared effects suggest that co-infection with these viruses could notably elevate the risk of AD development, as dysregulation of these processes are key pathological hallmarks of AD.

## Conclusions

In summary, our findings highlight the potential of these eight specific viruses to perturb crucial biological processes that are intricately linked to AD, reinforcing the connection between viral infections and the development and/or progression of AD. These perturbations can be mediated through shared and distinct mechanisms among viral species belonging to different categories, potentially contributing to variations in AD susceptibility based on the specific viral infection. Crucially, our findings suggest that reactivation of herpesviruses (HSV-1, HCMV and EBV) during acute infection with SARS-CoV-2 could potentially foster a detrimental interplay contributing to neurodegeneration. However, co-infection of SARS-CoV-2 with different pairs of herpesviruses leads to the amplification of dysregulation in different biological processes, contributing to variations in AD susceptibility risk. Nonetheless, the final outcome in AD susceptibility will be also influenced by various other risk factors, including genetic susceptibility, other environmental factors linked to AD, and the presence of comorbidities.

## Authors contribution

A.O. and P.Z have contributed to the conceptualization, review and editing of the manuscript. A.O. developed the methodology, collected and analyzed the data, and wrote the original draft of the manuscript. All authors have read and agreed to the published version of the manuscript.

## Data Availability Statement

This study utilized publicly available datasets for analysis. The data sources used are PHISTO (https://phisto.org/) and VirHostNet 3.0 (https://virhostnet.prabi.fr/).

## References

1. Deture MA, Dickson DW. The neuropathological diagnosis of Alzheimer’s disease. Mol. Neurodegener. 2019.

2. Folch J, Ettcheto M, Petrov D, et al. Review of the advances in treatment for Alzheimer disease: strategies for combating β-amyloid protein. Neurol (English Ed. 2018; 33(1):47–58.

3. World Health Organization. Dementia Fact sheet. Who. 2017. p. 1–4.

4. Migliore L, Coppedè F. Genetics, environmental factors and the emerging role of epigenetics in neurodegenerative diseases. Mutat. Res. - Fundam. Mol. Mech. Mutagen. 2009. p. 82–97.

5. Blackhurst BM, Funk KE. Viral pathogens increase risk of neurodegenerative disease. Nat. Rev. Neurol. 2023. p. 259–260.

6. Itzhaki RF, Dobson CB, Shipley SJ, Wozniak MA. The role of viruses and of APOE in dementia. Ann N Y Acad Sci. 2004. p. 15–18.

7. Lin WR, Shang D, Itzhaki RF. Neurotropic viruses and Alzheimer disease: Interaction of herpes simplex type I virus and apolipoprotein E in the etiology of the disease. Mol Chem Neuropathol. 1996; 28(1–3):135–141.

8. Wozniak MA, Shipley SJ, Combrinck M, Wilcock GK, Itzhaki RF. Productive herpes simplex virus in brain of elderly normal subjects and Alzheimer’s disease patients. J Med Virol. 2005; 75(2):300–306.

9. Moir RD, Lathe R, Tanzi RE. The antimicrobial protection hypothesis of Alzheimer’s disease. Alzheimer’s Dement. 2018; 14(12):1602–1614.

10. Zhou L, Miranda-Saksena M, Saksena NK. Viruses and neurodegeneration. Virol. J. 2013.

11. Sochocka M, Zwolińska K, Leszek J. The Infectious Etiology of Alzheimer’s Disease. Curr Neuropharmacol. 2017; 15(7).

12. Readhead B, Haure-Mirande JV, Funk CC, et al. Multiscale Analysis of Independent Alzheimer’s Cohorts Finds Disruption of Molecular, Genetic, and Clinical Networks by Human Herpesvirus. Neuron. 2018; 99(1):64–82.e7.

13. Carbone I, Lazzarotto T, Ianni M, et al. Herpes virus in alzheimer’s disease: Relation to progression of the disease. Neurobiol Aging. 2014; 35(1):122–129.

14. Kuhlmann I, Minihane AM, Huebbe P, Nebel A, Rimbach G. Apolipoprotein e genotype and hepatitis C, HIV and herpes simplex disease risk: A literature review. Lipids Health Dis. 2010.

15. Bortolotti D, Gentili V, Rotola A, Caselli E, Rizzo R. HHV-6A infection induces amyloid-beta expression and activation of microglial cells. Alzheimer’s Res Ther. 2019; 11(1).

16. Gold JE, Okyay RA, Licht WE, Hurley DJ. Investigation of long covid prevalence and its relationship to epstein-barr virus reactivation. Pathogens. 2021; 10(6).

17. Gatto I, Biagioni E, Coloretti I, et al. Cytomegalovirus blood reactivation in COVID-19 critically ill patients: risk factors and impact on mortality. Intensive Care Med. 2022; 48(6):706–713.

18. Seeßle J, Hippchen T, Schnitzler P, Gsenger J, Giese T, Merle U. High rate of HSV-1 reactivation in invasively ventilated COVID-19 patients: Immunological findings. PLoS One. 2021; 16(7 July).

19. Bond P. Ethnicity and the relationship between covid-19 and the herpes simplex viruses. Med. Hypotheses. 2021.

20. Hampshire A, Trender W, Chamberlain SR, et al. Cognitive deficits in people who have recovered from COVID-19. eClinicalMedicine. 2021; 39.

21. Becker JH, Lin JJ, Doernberg M, et al. Assessment of Cognitive Function in Patients after COVID-19 Infection. JAMA Netw Open. 2021; 4(10).

22. Matias-Guiu JA, Delgado-Alonso C, Yus M, et al. “Brain Fog” by COVID-19 or Alzheimer’s Disease? A Case Report. Front Psychol. 2021; 12.

23. Onisiforou A, Spyrou GM. Systems Bioinformatics Reveals Possible Relationship between COVID-19 and the Development of Neurological Diseases and Neuropsychiatric Disorders. Viruses [Internet]. 2022; 14(10):2270. Available from: 10.3390/v14102270

24. 2022 Alzheimer’s disease facts and figures. Alzheimer’s Dement. 2022; 18(4):700–789.

25. Durmus S, Çakir T, Özgür A, Guthke R. A review on computational systems biology of pathogen-host interactions. Front. Microbiol. 2015.

26. Khan MM, Ernst O, Manes NP, et al. Multi-omics strategies uncover host-pathogen interactions. ACS Infect. Dis. 2019. p. 493–505.

27. Onisiforou A, Spyrou GM. Immunomodulatory effects of microbiota-derived metabolites at the crossroad of neurodegenerative diseases and viral infection: network-based bioinformatics insights. Front Immunol. 2022; 13(843128).

28. Onisiforou A, Spyrou GM. Identification of viral-mediated pathogenic mechanisms in neurodegenerative diseases using network-based approaches. Brief Bioinform. 2021; 22(6).

29. Bindea G, Mlecnik B, Hackl H, et al. ClueGO: A Cytoscape plug-in to decipher functionally grouped gene ontology and pathway annotation networks. Bioinformatics. 2009; 25(8):1091–1093.

30. Tenenbaum D, Maintainer B. KEGGREST: Client-side REST access to the Kyoto Encyclopedia of Genes and Genomes (KEGG) [Internet]. 2022. p. R package version 1.38.0. Available from: https://rdrr.io/bioc/KEGGREST/

31. Kanehisa M, Goto S. KEGG: Kyoto Encyclopedia of Genes and Genomes. Nucleic Acids Res. 2000; 28(1):27–30.

32. Pemberton JR. Retention of mercurial preservatives in desiccated biological products. J Clin Microbiol. 1975; 2(6):549–551.

33. Durmuş Tekir S, Çakir T, Ardiç E, et al. PHISTO: Pathogen-host interaction search tool. Bioinformatics. 2013; 29(10):1357–1358.

34. Guirimand T, Delmotte S, Navratil V. VirHostNet 2.0: Surfing on the web of virus/host molecular interactions data. Nucleic Acids Res. 2015; 43(D1):D583–D587.

35. Meysman P, Titeca K, Eyckerman S, et al. Protein complex analysis: From raw protein lists to protein interaction networks. Mass Spectrom. Rev. 2017. p. 600–614.

36. Huang SY, Yang YX, Kuo K, et al. Herpesvirus infections and Alzheimer’s disease: a Mendelian randomization study. Alzheimer’s Res Ther. 2021; 13(1).

37. Barnes LL, Capuano AW, Aiello AE, et al. Cytomegalovirus infection and risk of alzheimer disease in older black and white individuals. J Infect Dis. 2015; 211(2):230–237.

38. Manoharan S, Guillemin GJ, Abiramasundari RS, Essa MM, Akbar M, Akbar MD. The Role of Reactive Oxygen Species in the Pathogenesis of Alzheimer’s Disease, Parkinson’s Disease, and Huntington’s Disease: A Mini Review. Oxid. Med. Cell. Longev. 2016.

39. Uddin MS, Stachowiak A, Mamun A Al, et al. Autophagy and Alzheimer’s disease: From molecular mechanisms to therapeutic implications. Front. Aging Neurosci. 2018.

40. Tiwari S, Atluri V, Kaushik A, Yndart A, Nair M. Alzheimer’s disease: Pathogenesis, diagnostics, and therapeutics. Int. J. Nanomedicine. 2019. p. 5541–5554.

41. Patten DA, Germain M, Kelly MA, Slack RS. Reactive oxygen species: Stuck in the middle of neurodegeneration. J. Alzheimer’s Dis. 2010.

42. Menzies FM, Fleming A, Caricasole A, et al. Autophagy and Neurodegeneration: Pathogenic Mechanisms and Therapeutic Opportunities. Neuron. 2017.

43. Eimer WA, Vijaya Kumar DK, Navalpur Shanmugam NK, et al. Alzheimer’s Disease-Associated β-Amyloid Is Rapidly Seeded by Herpesviridae to Protect against Brain Infection. Neuron. 2018; 99(1):56–63.e3.

44. Vossen MTM, Westerhout EM, Söderberg-Nauclér C, Wiertz EJHJ. Viral immune evasion: A masterpiece of evolution. Immunogenetics. 2002; 54(8):527–542.

45. Choi HG, Soh JS, Lim JS, Sim SY, Lee SW. Association between dementia and hepatitis B and C virus infection. Med (United States). 2021; 100(29):E26476.

46. Huang L, Wang Y, Tang Y, He Y, Han Z. Lack of Causal Relationships Between Chronic Hepatitis C Virus Infection and Alzheimer’s Disease. Front Genet. 2022; 13.

47. Tran L, Jung J, Carlin C, Lee S, Zhao C, Feldman R. Use of Direct-Acting Antiviral Agents and Survival among Medicare Beneficiaries with Dementia and Chronic Hepatitis C. J Alzheimer’s Dis. 2021; 79(1):71–83.

48. Chatterjee D, Biswas K, Nag S, Ramachandra SG, Sarma J Das. Microglia play a major role in direct viral-induced demyelination. Clin Dev Immunol. 2013; 2013.

49. Pflugrad H, Meyer GJ, Dirks M, et al. Cerebral microglia activation in hepatitis C virus infection correlates to cognitive dysfunction. J Viral Hepat. 2016; 23(5):348–357.

50. Tekir SD, Çakir T, Ülgen KÖ. Infection strategies of bacterial and viral pathogens through pathogen-human protein-protein interactions. Front Microbiol. 2012; 3(FEB).

51. Dyer MD, Murali TM, Sobral BW. The landscape of human proteins interacting with viruses and other pathogens. PLoS Pathog. 2008; 4(2).

52. DaPalma T, Doonan BP, Trager NM, Kasman LM. A systematic approach to virus-virus interactions. Virus Res. 2010. p. 1–9.

53. Lavadi RS, Orrick JA. Biochemistry, Protein Targeting and I Cell Diseases. StatPearls. 2023; .

54. Talebi M, Mohammadi Vadoud SA, Haratian A, et al. The interplay between oxidative stress and autophagy: focus on the development of neurological diseases. Behav. Brain Funct. 2022.

