## Supplementary File 1 for "From Viral Infections to Alzheimer’s Disease: Unveiling the Mechanistic Links Through Systems Bioinformatics"

**Supplementary Table 1**: Characteristics of the integrated Virus-Host-AD-PPI networks.

| | Nodes | Edges | Number of proteins in the AD subnetwork | Number of proteins in the virus subnetwork |
|---|---|---|---|---|
| HSV1-host-AD PPI network | 880 | 7522 | 459 | 759 |
| EBV-host-AD PPI network | 1590 | 13001 | 667 | 1407 |
| HCV-host-AD PPI network | 1355 | 6386 | 584 | 1153 |
| HCMV-host-AD PPI network | 2387 | 13458 | 871 | 2187 |
| SARS-CoV2-host-AD PPI network | 2914 | 25851 | 1036 | 2874 |
| HBV-host-AD PPI network | 353 | 1526 | 263 | 199 |
| KSHV-host-AD PPI network | 1066 | 5360 | 465 | 878 |
| IAV-host-AD PPI network | 3184 | 11887 | 2832 | 2800 |

**Supplementary Table 2**: Number of GO biological processes found to be associated with the extracted subnetworks from the eight integrated Virus-Host-AD PPI networks, as well as the overlapping GO biological processes (Viruses ∩ AD).

| | AD subnetwork GO Biological processes | Virus subnetwork GO Biological processes | Overlapping GO Biological processes (Viruses ∩ AD) |
|---|---|---|---|
| HSV1-host-AD PPI network | 941 | 620 | 559 |
| EBV-host-AD PPI network | 958 | 587 | 561 |
| HCV-host-AD PPI network | 882 | 479 | 497 |

| | | | |
|---|---|---|---|
| HCMV-host-AD PPI network | 1001 | 421 | 399 |
| SARS-CoV2-host-AD PPI network | 981 | 418 | 303 |
| HBV-host-AD PPI network | 988 | 672 | 591 |
| KSHV-host-AD PPI network | 1098 | 633 | 570 |
| IAV-host-AD PPI network | 485 | 380 | 363 |
